# CellWalkR: An R Package for integrating single-cell and bulk data to resolve regulatory elements

**DOI:** 10.1101/2021.02.23.432593

**Authors:** Pawel F. Przytycki, Katherine S. Pollard

## Abstract

While single-cell open chromatin (scATAC-seq) data allows for the identification of cell type-specific regulatory regions, it is much sparser than bulk data. CellWalkR is an R package that performs an integration of external labeling and bulk epigenetic data with scATAC-seq using a network-based random walk model to help overcome this sparsity. Outputs include cell type labels for individual cells and regulatory regions.

**Availability and implementation:** CellWalkR is freely available as an R package under a GNU GPL-2.0 License, and can be accessed from https://github.com/PFPrzytycki/CellWalkR with an accompanying vignette for analyzing example data.

## Introduction

Gene regulatory elements active in specific cell types can be identified using scATAC-seq. However, this data is notoriously low-resolution with high rates of dropout, making analysis difficult^1^. Bulk data has much higher resolution which allows for high confidence identification of regulatory regions, but does not provide cell type-specific annotations. Single-cell RNA sequencing (scRNA-seq) is less sparse than scATAC-seq and hence can identify dozens of cell types in complex tissues, but it does not pinpoint regulatory elements^1^.

We developed CellWalkR, an R package that combines the strengths of these different technologies to resolve regulatory elements to cell types (Figure 1a). The package implements and extends a previously introduced network-based model^2^ that relies on a random walk with restarts model of diffusion. Additional features include filter inputs, flexible scATAC-seq formats, multiple label sets, and a GPU implementation using TensorFlow^3^. The output is a large influence matrix, portions of which are used for cell labeling, determining label similarity, embedding cells into low dimensional space, and mapping regulatory regions to cell types.

**Figure 1.**
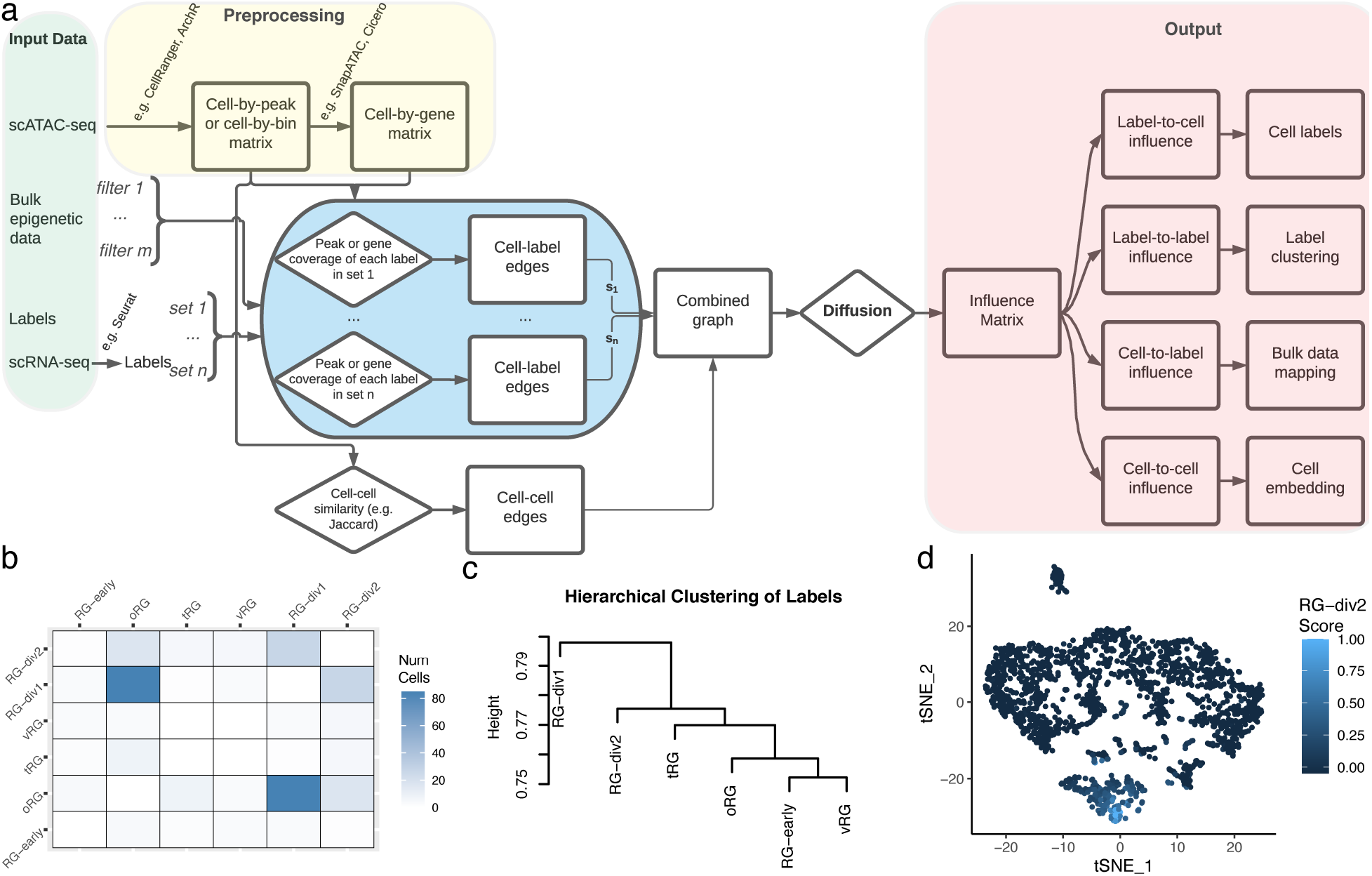
Overview of CellWalkR. **a**. Flowchart of CellWalkR pipeline. CellWalkR takes scATAC-seq, bulk epigenetic data, and label sets as input which it then combines into a graph. After diffusing information across this graph, different portions of the resulting influence matrix are used for analysis. **b**. Uncertainty matrix showing how many cells have nearly the same influence from two different labels. This example uses label data for radial glia based on scRNA-seq^9^ and scATAC-seq from a population of radial glia in the developing brain^10^. **c**. Hierarchical clustering of labels based on label-to-label influence **d**. t-SNE embedding of cells based on cell-to-cell influence with the label influence for one cell type shown. Abbreviations: Radial Glia (RG), outer RG (oRG), truncated RG (tRG), and ventral RG (vRG).

## Results

CellWalkR builds a network consisting of two types of nodes representing cells and labels (i.e., cell types or states). Building this network requires two types of input: a scATAC-seq experiment and labeling data. Users can input scATAC-seq data as a cell-by-region matrix (e.g. cell-by-peak or cell-by-bin) or as processed by SnapATAC^4^, ArchR^5^, or Cicero^6^. This input is used to compute the similarity between cells, using Jaccard similarity by default, though any function can be used. Cell similarity gives the weights on cell-cell edges.

In order to generate edges between labels and cells CellWalkR links genomic regions in scATAC-seq to cell types or states through marker genes. An edge is generated between a label and a cell based on how open the marker genes are in the cell’s scATAC-seq data. The definition of regions used to quantify openness of marker genes is flexible and user-defined; it can correspond to promoters, gene bodies, and/or distal regions (e.g. correlated peaks identified by Cicero). The user also provides a file that associates each label with a set of gene names and optionally their log-fold changes in expression for that label. CellWalkR includes a function to generate this from scRNA-seq data using Seurat^7^ if desired. This strategy can be used to build networks for an arbitrary number of label sets, representing cell types in different time points or disease states.

All label-cell and cell-cell edges are combined into a single network, with each set of labeling edges scaled by an associated parameter *s*. Once a complete network is built, a random walk with restarts model of diffusion is run to convergence^2^. CellWalkR can optionally run this step on a GPU using TensorFlow^3^ for a greater than 15-fold speedup (Supplementary Figure 1a). With this speedup, CellWalkR can be run on 10,000 cells in under 10 seconds using 12Gb of memory (Supplementary Figure 1b). The output of this process is an influence matrix where each column represents the amount of information that flows from each node to each other node.

In addition to labeling data, CellWalkR can also take bulk epigenetic data as input. This data serves as a filter through which scATAC-seq peaks are passed before being considered for calculating cell-label edges. Each filter has a scaling parameter *f* for each label set that determines the strength of the filter. Filters are restrictive by default (remove scATAC-seq peaks overlapping filter regions) but can also be set to permissive (allow only overlapping peaks). They can apply to the entire associated gene or just overlapping peaks.

Setting the *s* parameter and the *f* parameters correctly for each label set can make a large difference for how information is propagated through the network. CellWalkR has built-in functions that tune these parameters to optimize cell homogeneity, a measure of influence between cells of the same label versus between cells of different labels^2^. To accelerate optimization, CellWalkR can downsample cells and can run these calculations in parallel.

CellWalkR has many options for downstream analysis, all of which stem from using different portions of the influence matrix. The label-to-cell influence portion is used to assign labels to cells. These label estimates are fuzzy, meaning each cell has a distribution of scores from each label. Thanks to this, CellWalkR can determine how often cells receive nearly the same score for two different labels, which it summarizes in an uncertainty matrix (Figure 1b). For deterministic labeling, CellWalkR assigns the label with maximum influence. CellWalkR can also estimate how similar labels are to each other using label-to-label influence, and these similarities can be used for hierarchical clustering of cell types or other labels (Figure 2b).

Visualizing cells in a two-dimensional space is a helpful strategy for understanding how cells relate to each other and to their labels. CellWalkR uses t-SNE^8^ to embed cell-to-cell influence and has several options for coloring cells. First, they can be colored by their assigned label, optionally marking cells as unknown if the maximum influence on the cell is below a threshold (Supplemental Figure 2a, left). Embedding cell-to-cell influence rather than sparse, noisy cell accessibility profiles has clear advantages for separating cells of different types, especially rare cell types (Supplemental Figure 2a and 2b). CellWalkR can also generate t-SNE plots showing the amount of influence a single label has on each cell (Figure 1d) or the difference in influence between two labels, allowing the user to identify transition regions.

Many genomic regions are annotated by bulk data without cell type-specific annotations. Examples include genetic variants, predicted enhancers, transcription factor binding sites, and epigenetic segmentations. CellWalkR can associate such regions to cell types using cell-to-label influence. For each region provided by the user, CellWalkR calculates its influences on each label. In this way any type of genomic region can be mapped to labels, as long as cells of the appropriate type or state are in the scATAC-seq data.

## Conclusions

Many regulatory elements are only mapped in bulk data, and scATAC-seq data can be difficult to use due to its sparsity. CellWalkR is a flexible R package for multi-label analysis of scATAC-seq data incorporating bulk epigenetic and expression data to label cells. The package can resolve rare cell types and associate regulatory elements with those cell types.

## Acknowledgements

Thank you to the many people who tested CellWalkR including Sean Whalen, Amanda Everitt, Candace Chan, Maureen Pittman, Patrick Bradley, and Laura Gunsalus.

## Funding

This work was supported by research grants to K.S.P from: NIH/NIMH awards U01-MH116438, R01-MH109907, and R01-MH123178. Funding bodies played no role in the design of the study and collection, analysis, and interpretation of data, or in writing the manuscript.

## Notes

### Competing Interest Statement

The authors have declared no competing interest.

