## Supplemental Figures for "CellWalkR: An R Package for integrating single-cell and bulk data to resolve regulatory elements"

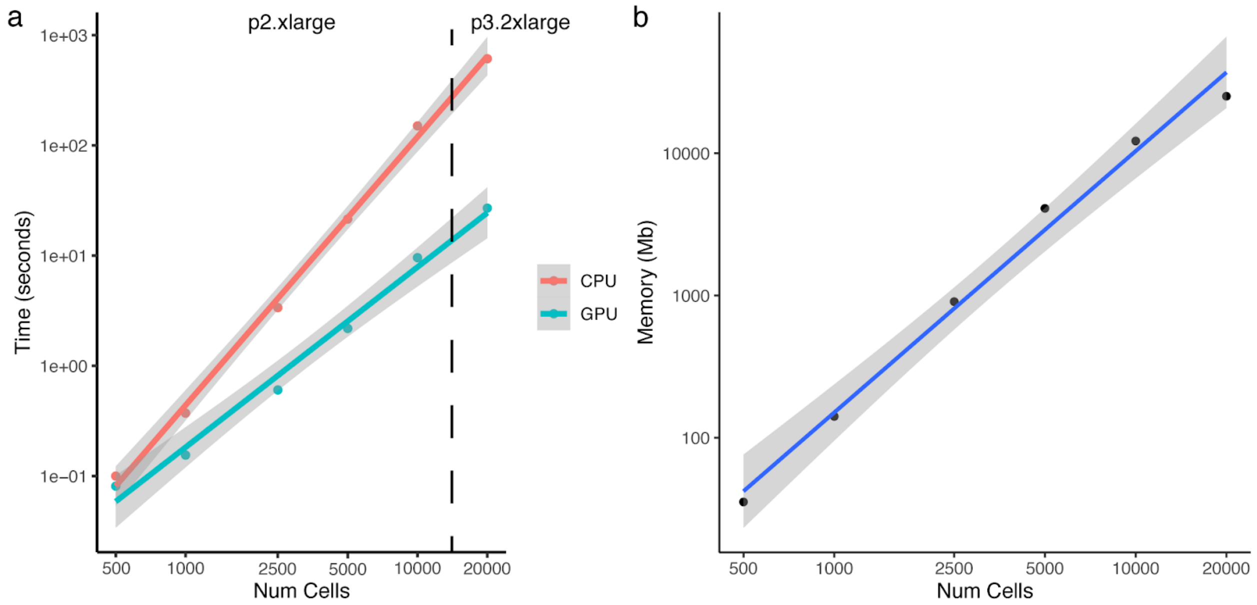


Supplemental Figure 1. **CellWalkR Performance. a.** Run time of CellWalkR on AWS P2.xlarge instance (for up to 15,000 cells) and P3.2xlarge (for 20,000 cells) using a CPU or GPU. The GPU is more than 15 times faster, with greater benefits for more cells. **b.** Memory used by CellWalkR on a P2.xlarge instance.

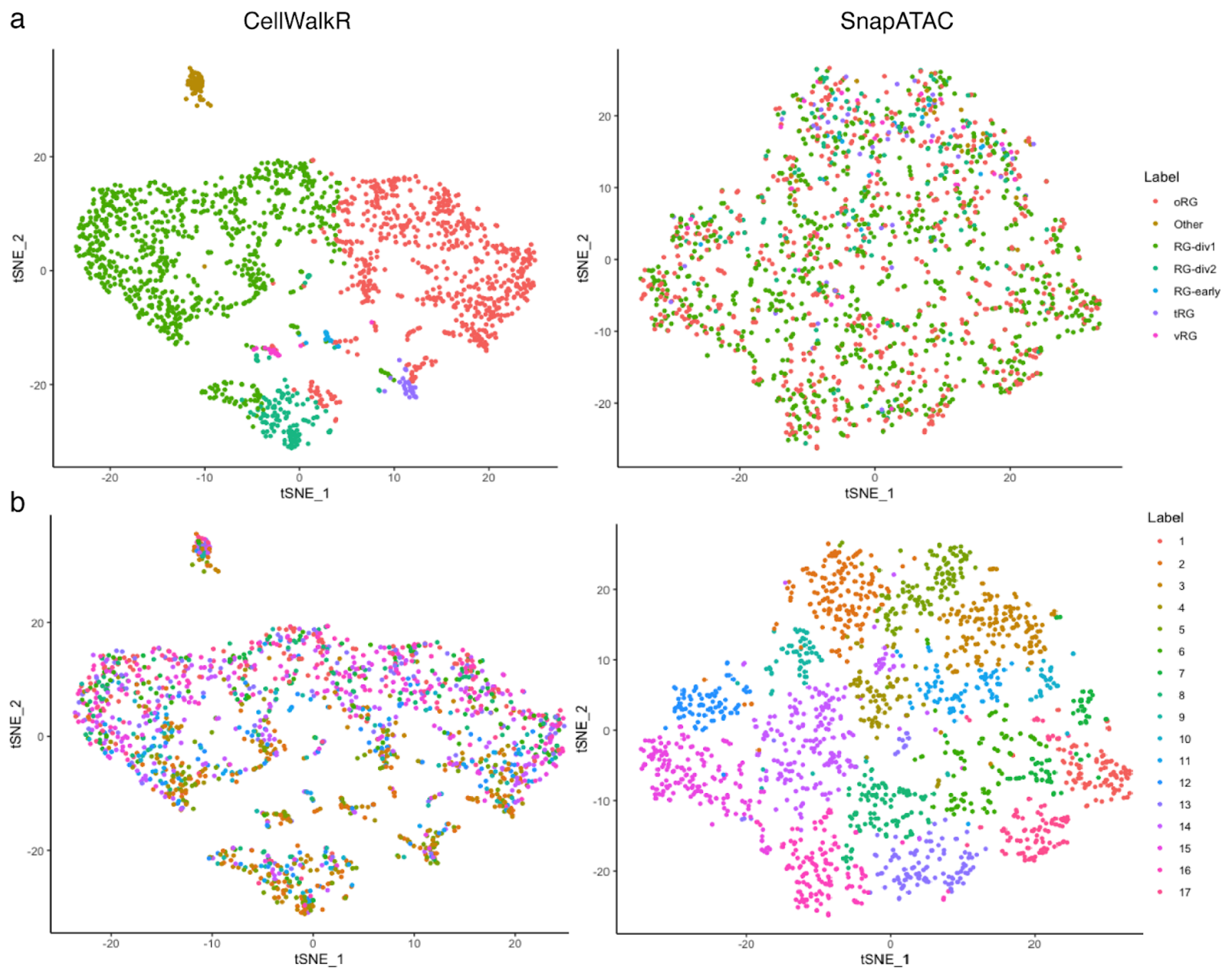


Supplemental Figure 2. **Comparison to SnapATAC. a.** CellWalkR embedding of cell-to-cell influence using t-SNE (left) creates more distinct clusters than the embedding generated by SnapATAC (right) with clear separation of labeled cell types. Cells are colored by maximally influencing labels in CellWalkR, with cells received a maximum influence less than 0 marked as "Other." Note that labels are not defined by clustering in embedding, thus the grouping of cell types in the embedding can serve as a validation of the distinctness of labels. Additionally, CellWalkR displays an ability to identify both common cell types and very rare cell types, with a large dynamic range in cluster size. **b.** Using the same embeddings as in panel a, but now with cells colored by their cluster assignment according to SnapATAC. SnapATAC detects a very large number of clusters and has no built in ability to detect what cell type they represent. The clusters all include similar numbers of cells (right). When these cluster assignments are plotted in the CellWalkR embedding (left), there is a clear gradient from top to bottom but no separation between labels.
